# Characterizing scaling laws in gut microbial dynamics from time series data: caution is warranted

**DOI:** 10.1101/2021.01.11.426045

**Authors:** Xu-Wen Wang, Yang-Yu Liu

## Abstract

Many studies have revealed that both host and environmental factors can impact the gut microbial compositions, implying that the gut microbiota is considerably dynamic^1–5^. In their Article, Ji et al.^6^ performed comprehensive analysis of multiple high-resolution time series data of human and mouse gut microbiota. They found that both human and mouse gut microbiota dynamics can be characterized by several robust scaling laws describing short- and long-term changes in gut microbiota abundances, distributions of species residence and return times, and the correlation between the mean and the temporal variance of species abundances. They claimed that those scaling laws characterize both short- and long-term dynamics of gut microbiota. However, we are concerned that their interpretation is quite misleading, because all the scaling laws can be reproduced by the shuffled time series with completely randomized time stamps of the microbiome samples.

## Introduction

Let us consider a time series of gut microbial compositions {*X*_*k*_(*t*)}, where *X*_*k*_(*t*) represents the relative abundance of OTU-*k*, *k* = 1, ··· *N*, and *t* = 1, ··· *T*. Ji et al. proposed multiple scaling laws to describe the dynamics of gut microbiota (**Fig.1**, solid dots):

1. Short-term (daily) abundance change *μ* follows a Laplace distribution: 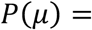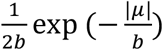, which has a characteristic tent shape in log-transformed probabilities. Here, *μ* is calculated by averaging *μ*_*k*_(*t*) ≡ log(*X*_*k*_(*t* + 1)/*X*_*k*_(*t*)) across all OTUs and all time points, and *b* is a scale parameter.
2. The variability (i.e., standard deviation) of daily abundance change *σ*_*μ*_ decreases approximately linearly with the increasing mean of successive log-abundance *x*_*m*_(*t*), i.e., *σ*_*μ*_ = *rx*_*m*_ + *c*, where 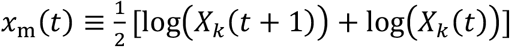, *r* is the slope, and *c* is a constant.
3. The long-term drift of gut microbiota abundance can be approximated by the equation of anomalous diffusion: 〈*δ*^2^(Δ*t*)〉 ∝ Δ*t*^2*H*^, where 〈*δ*^2^(Δ*t*)〉 is calculated by averaging 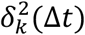 across all OTUs and all time points, *δ*_*k*_(Δ*t*) ≡ log(*X*_*k*_(*t* + Δ*t*)/*X*_*k*_(*t*)), and *H* is the Hurst exponent.
4. The distribution of residence time *t*_res_ (or return times *t*_ret_) follows a power law with exponential tail: *P*(*t*) ∝ *t*^−*α*^e^−*λt*^. Here, *t*_res_ (or *t*_ret_) is the time interval during which an OTU was continuously detected (or absent) from the community, respectively. The exponential tail e^−*λt*^ results from the finite length of the analyzed time series, and *α* is the power-law exponent.
5. The temporal variability patterns of gut microbiota closely follow Taylor’s law: 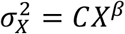, where *C* is a constant, *X* and 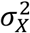 are the mean and variance of species abundance, *β* is the power-law exponent.

**Figure 1:**
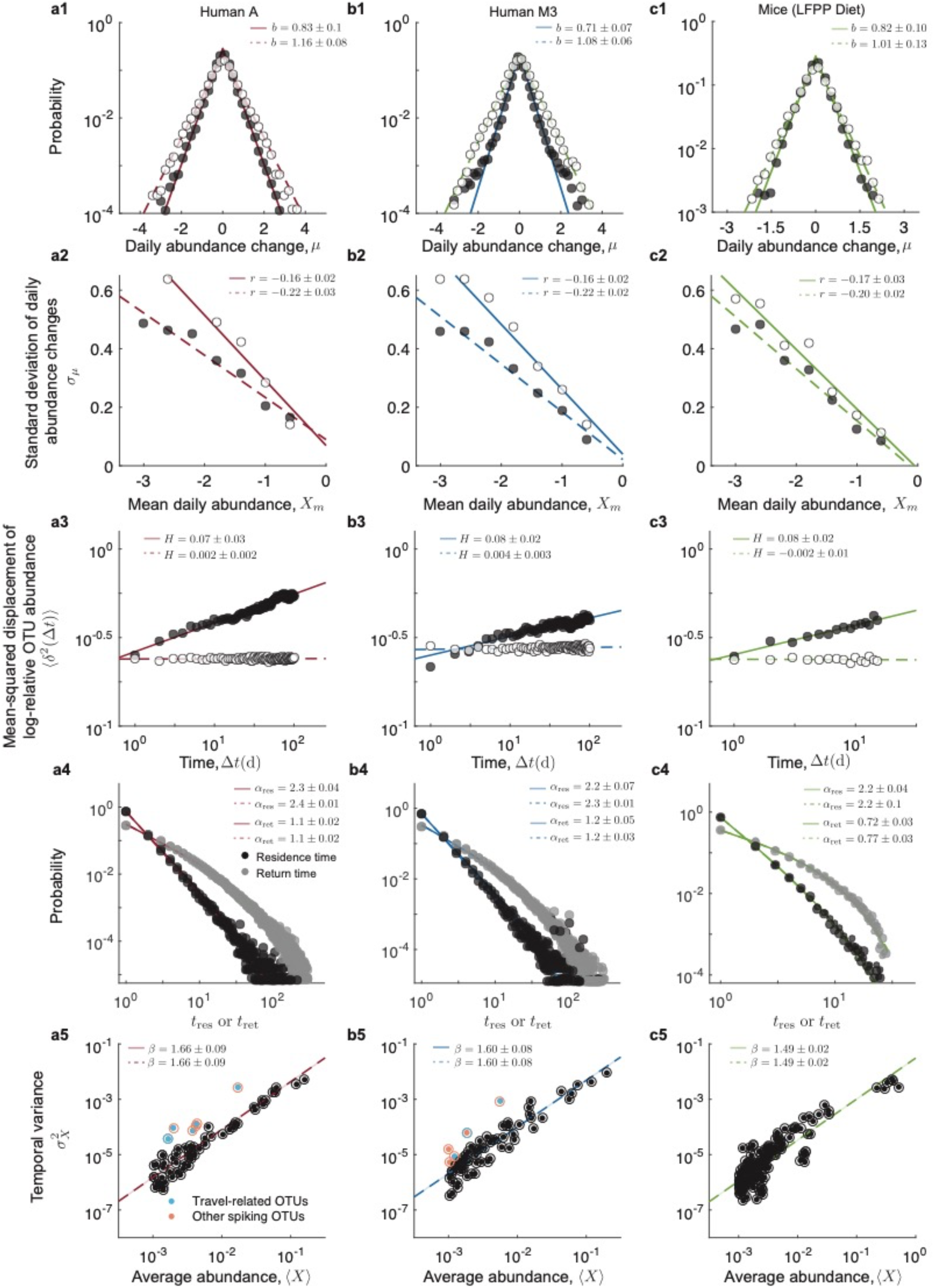
Scaling relationships observed from the time series data of human and mouse gut microbiota can also be observed from the shuffled time series. Throughout this figure, solid (or hollow) dots represent results obtained from the original (or shuffled) time series, respectively. Lines represent maximum likelihood estimation (MLE) fits to the data. To control for known technical factors such as sample preparation and sequencing noise, here we adopted exactly the same OTU inclusion criteria as used by Ji et al^6^. **Columns:** (**a**) Human A; (**b**) Human M3; (**c**) Mice fed on a low-fat, plant polysaccharide (LFPP) diet. **Rows:** (**1)**Short-term (daily) abundance change *μ* follows a Laplace distribution: 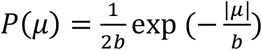. (**2)** The variability (i.e., standard deviation) of daily abundance change *σ*_*μ*_ decreases approximately linearly with increasing the mean of successive log-abundance *x*_*m*_(*t*). (**3**) The long-term drift of gut microbiota abundance can be approximated by the equation of anomalous: 〈*δ*^2^(Δ*t*)〉 ∝ Δ*t*^2*H*^, where *H* is the so-called Hurst exponent. (**4**) The distributions of residence (*t*_res_) and return times (*t*_ret_) follow power laws with exponential tails: *P*(*t*) ∝ *t*^−*α*^e^−*λt*^. (**5**) The temporal variability patterns of gut microbiota closely follow Taylor’s law: 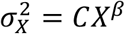, where *C* is a constant, *X* and 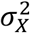 are the mean and variance of species abundance.

## Results

We emphasize that, without comparing with appropriate null models^7^, those scaling laws founded by Ji et al. may not reflect the true complex dynamics of gut microbiota. Indeed, after shuffling the time series (i.e., randomly relabeling the time stamp of each microbiome sample), we can recover all the scaling laws up to the change of a few exponent values (**Fig.1**, hollow dots). For certain scaling laws (e.g., the power-law distributions of *t*_res_ and *t*_ret_), the shuffling will even keep the power-law exponents almost unchanged (**Fig.1**, Row-4). As for Taylor’s law, it will not be affected by the shuffling at all (**Fig.1**, Row-5).

The shuffled time series represents a very natural null model, where the temporal structure in the original time series has been largely destroyed. Given that such a null model with completely randomized time stamps can reproduce all the scaling laws, those scaling laws observed from the real time series data of human and mouse gut microbiota should not be simply characterized as short-term or long-term dynamics of gut microbiota.

To better understand the stochasticity in time series data of human and mouse gut microbiota, and quantify their temporal structure, we computed the noise-type profile^8^ for each time series analyzed by Ji et al. Here, the noise type of an OTU is obtained by decomposing its relative abundance fluctuations into spectral densities at specific frequencies using a Fourier transform. The slope in the spectral density vs. frequency log-log plot is used to distinguish black (below −2), brown (around −2), pink (around −1), and white noise (no negative slope). The dependency on previous time points is strongest for black noise, weakest for pink noise, and is absent for white noise. We found that the noise type is dominated by white and pink noises for all the gut microbiota time series analyzed by Ji et al., indicating the very weak temporal structure in those time series (**Fig.2**, ‘Real’). Note that the shuffled time series (our null model) displays a much higher percentage of white noise, indicating that the weak temporal structure in the original time series has been further destroyed (**Fig.2**, ‘Null’).

**Figure 2:**
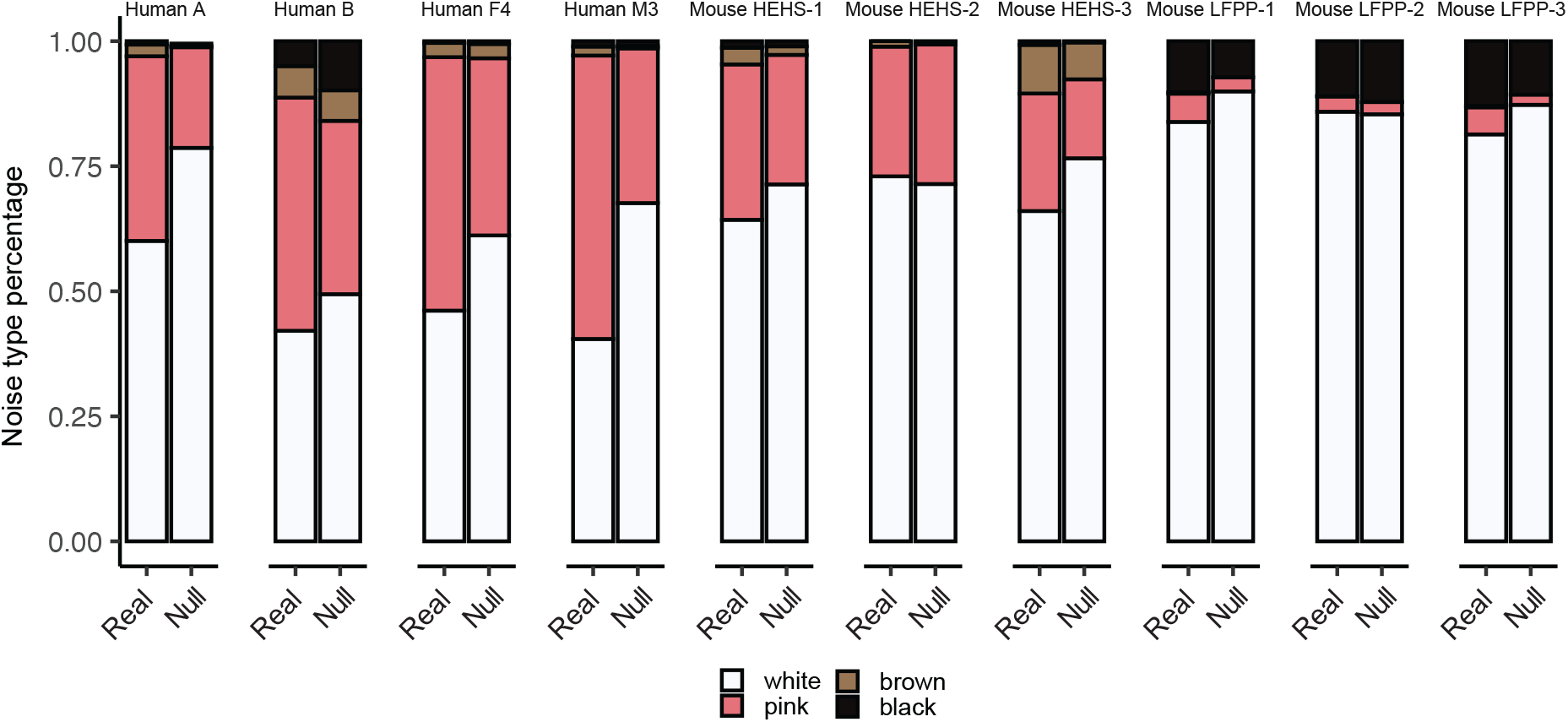
Noise-type profiles of human and mouse gut microbiota time series. The bar plots depict for each host’s gut microbiota time series the percentage of OTUs with white, pink, brown, or black noise. White noise indicates the absence of temporal structure, all other noise types indicate the presence of certain levels of temporal structure: the dependency on previous time points is strongest for black noise, medium for brown noise, and weakest for pink noise. For the noise-type profile analysis, we interpolated the data with function “stineman” in the R package stinepack^10^ to ensure equidistant time intervals. Also, the last time point from Human B was omitted, since there was a gap of 66 days between it and the previous sample. Labels: ‘Real’ represents the original time series, while ‘Null’ represents the shuffled time series. HEHS: high-fat, high-sugar. LFPP: low-fat, plant polysaccharide.

Our results are consistent with previous findings that fluctuations in microbial abundance are mainly due to temporal stochasticity^9^, and human gut microbiota has two distinct dynamic regimes: auto-regressive and non-autoregressive^1^. In particular, most of the variance in gut microbial time series is non-autoregressive and driven by external day-to-day fluctuations in host and environmental factors (e.g., diet), with occasional internal autoregressive dynamics as the system recovered from larger shocks (e.g. facultative anaerobe blooms)^1^. Overall, the gut microbiota can be considered as a dynamically stable system, continually buffeted by internal and external forces and recovering back toward a conserved steady-state^1^. This picture also qualitatively explains why the shuffled time series yields the same scaling laws as the original time series does. Both the original and shuffled time series of gut microbiota have very weak temporal structure. Hence, generally speaking, data analysis that explicitly considers the time stamps of microbiome samples will yield very similar results.

Taken together, the scaling laws observed by Ji et al. should not be interpreted as the characteristics of short-term or long-term dynamics of gut microbiota. They do not represent the autoregressive or internal ecological dynamics of gut microbiota but represent the dominating non-autoregressive dynamics of gut microbiota driven by external environmental fluctuations. The exact mechanisms generating these scaling laws certainly warrant further investigation.

## Author Contributions

Y.-Y.L. conceived and designed the project. X.-W.W. performed all the numerical calculations and data analysis. Both authors interpreted the results and wrote the manuscript.

## Author Information

The authors declare no competing financial interests. Correspondence and requests for materials should be addressed to Y.-Y.L.

